# Label-free imaging of matrix mineralization in alginate-encapsulated bone spheroids using Coherent Raman Scattering microscopy

**DOI:** 10.64898/2026.01.27.701971

**Authors:** Diamante Boscaro, Sebastian Kjeldgaard-Nintemann, Astrid Bjørkøy, Pawel Sikorski

## Abstract

Three-dimensional (3D) cell cultures, such as spheroids, are increasingly used to perform advanced studies on bone matrix mineralization. However, their full characterization remains challenging. Traditional colorimetric and fluorescent assays using dyes, such as Alizarin Red S (ARS) and calcein, are effective in monolayer cell cultures, but fail to provide reliable information when used in complex 3D cell constructs. In this study, we investigated the application of Coherent Raman Scattering microscopy for label-free, comprehensive characterization of extracellular matrix (ECM) mineralization in alginate-encapsulated bone spheroids. After confirming that traditional staining techniques are unreliable for mineral detection in spheroids, Stimulated Raman Scattering (SRS) microscopy was used to detect phosphate-rich mineral deposits at a Raman shift of 960 cm^−1^, while Second Harmonic Generation (SHG) microscopy was used in association with SRS to provide complementary information on the deposition and organization of the collagenous matrix. SRS was used to detect lipid-rich regions at a Raman shift of 2857 cm^−1^ to perform cell localization. SRS imaging revealed the presence of phosphate-rich regions in the spheroids, including the core regions, usually challenging to characterize in intact 3D constructs. Raman spectral scans on SRS-positive regions confirmed the specificity of the phosphate signal. In addition, comparison of SRS and Coherent Anti-Stokes Raman Scattering (CARS) demonstrated the advantage of SRS in terms of reduced background compared to CARS for lipid imaging. Taken together, our results demonstrated that SRS, in combination with SHG, provides a promising and powerful approach to perform label-free, chemically specific characterization of intact 3D bone models.

## 1 Introduction

In bone tissue engineering (BTE), accurate assessment of mineral deposition is crucial for evaluating the osteogenic potential of *in vitro* bone cell models. Bone is a hierarchically organized tissue, composed of cells and mineralized extracellular matrix (ECM) [1]. Bone ECM is composed of an organic part (40%), consisting mainly of type I collagen (90%), and an inorganic part (60%), consisting of calcium-deficient apatite [2–4]. The precise relationship between the organic and inorganic components is essential for the correct mechanical and biological functions of bone [4] and for this reason, proper evaluation of these components is an essential aspect in BTE studies. One of the more common cell lines used for investigating bone mineralization is the murine cell line MC3T3-E1, as the cells are easy to maintain and differentiation can be induced by treatment with ascorbic acid and *β*-glycerophosphate [5–8].

Traditionally, two-dimensional (2D) cell cultures have been used as the standard model for understanding the process of bone mineralization [9]. 2D cell cultures, however, fail to replicate specific aspects of the natural bone microenvironment, such as the extensive cell-cell and cell-ECM interactions [10]. In response to these limitations, three-dimensional (3D) cell models, such as spheroids, have been developed [11]. 3D cell models have the advantage of replicating the natural microenvironment of the tissue of interest, thereby offering a more relevant cell model for *in vitro* studies [12]. In particular, spheroids have been shown to replicate physiological cell-cell and cell-ECM interactions [13], making them an optimal model for complex *in vitro* bone studies.

Traditional methods for assessing bone mineralization in *in vitro* cell models are based on colorimetric assays, such as Alizarin Red stain (ARS) or calcein labeling, or more advanced methods such as electron microscopy. Alizarin Red S is a salt that chelates calcium deposits, staining them with red color [14], while calcein is a fluorescent calcium-binding molecule [15]. While these methods have been extensively used for the detection of calcium phosphate (CaP) deposits, they are not specific for calcium in CaP deposits, resulting in the possibility of staining other calcium sources in the sample [6].

The increased application of spheroids in BTE studies resulted in the need of analytical methods that allow complete characterization of these cell models[16]. Traditional colorimetric or imaging techniques, which have been developed and optimized for 2D cell models, can face challenges when applied to 3D cell cultures, mostly as a result of the thickness of the sample [17]. These limitations often make extensive sample handling necessary, which can lead to sample loss, damage or introduction of artifacts.

One of the possible approaches to address these issues is the application of label-free methods that can assess the biochemical composition of the sample, while being minimally invasive or requiring minimal sample handling. Raman spectroscopy and microscopy are label-free techniques used to assess the molecular composition of biological samples by probing molecular vibrations. Raman spectroscopy is based on the inelastic scattering of photons, in which a photon interacts with matter leading to the exchange of energy and a change of the vibrational energy state within the molecular bonds of the participating molecule. The amount of energy exchanged (and thus the change in wavelength of the Raman-scattered photons) can give information about the molecular bonds present in the sample [18, 19]. In Raman microscopy, imaging of Raman-scattered photons arising from a specific energy transition allows for non-destructive and non-invasive chemical imaging, while at the same time allowing for imaging of living cells [20]. For bone mineralization studies, Raman spectroscopy and microscopy have proven to be effective in the detection of CaP deposits in monolayer cell cultures [21, 22], cells grown on a scaffold [23] and bone tissue sections [24, 25]. On the other hand, only a few studies have reported the application of Raman spectroscopy and microscopy on intact 3D cell models, while others perform sectioning of the spheroids prior to analysis [26–29]. It is important to notice that while Raman spectroscopy is a valuable method for biochemical characterization of biological samples, it is characterized by a relatively weak signal, in addition to the Raman signal being possibly influenced by the presence of fluorescent molecules in biological samples [30].

In this context, Coherent Raman Scattering (CRS) microscopy can be used to overcome the limitations of spontaneous Raman Scattering. CRS microscopy comprises two main methods: Coherent Anti-Stokes Raman Scattering (CARS) and Stimulated Raman Scattering (SRS) microscopy. Both of these techniques are non-linear, label-free and non-invasive methods that use two laser beams, called pump and Stokes beam, respectively, to stimulate vibrational energy transitions within the molecular bonds present in the sample. These energy transitions can be probed by tuning the energy difference between the two beams to match a known Raman shift, and thereby, CRS allows for chemical characterization of the sample in much the same way as spontaneous Raman microscopy [31, 32]. In both CARS and SRS, the intensity of the Raman signal is increased by a few orders of magnitude, compared to spontaneous Raman scattering[33].

In CARS microscopy, the two laser beams are used to facilitate the transition of a molecular bond to a higher vibrational energy level and then back to the initial state, resulting in the release of an anti-Stokes photon. This complex process can be described as a four-wave mixing process [34]. Even though CARS is characterized by a stronger signal compared to spontaneous Raman scattering, a big disadvantage for its application, especially for studies on biological samples, is the limited contrast due to the non-resonant background [35, 36].

In SRS microscopy, the frequency difference between the pump and the Stokes beam is set to match the vibrational frequency of a molecule, resulting in coherent excitation of the corresponding chemical bond [33, 37]. As a result, an intensity loss is observed in the pump beam (Stimulated Raman Loss) and an intensity gain is observed in the Stokes beam (Stimulated Raman Gain)[38]. These intensity changes are measured and used to generate an SRS image [39]. SRS can be used to analyze the chemical composition of the sample and evaluate the concentration of the components of interest, as the SRS signal is directly correlated to the concentration of the targeted chemical groups [33]. In addition, SRS is a filter-free technique, as the signal is measured as a change in the intensity of the the pump or Stokes beam [39]. Thus, contrary to CARS, in SRS microscopy there is no non-resonant background, resulting in SRS spectra that closely correspond to those of spontaneous Raman scattering [33, 37]. Because CRS is a multi-photon technique, like Second Harmonic Generation (SHG) microscopy, it benefits from the advantages observed with multi-photon imaging, such as increased imaging depth and great optical sectioning without the need for a pinhole. These features make CRS a well-suited method for imaging 3D cell models.

We have previously investigated the application of alginate-encapsulated bone spheroids as a model to study bone mineralization. In particular, MC3T3-E1 spheroids were characterized and SHG microscopy was used to perform label-free imaging of the deposited collagenous matrix, demonstrating the advantage of this technique in providing comprehensive information about collagen deposition in the intact, 3D structures [40]. In this work, we evaluated the applicability of traditional colorimetric and fluorescent assays, such as ARS, calcein staining and OsteoImage™, for detecting CaP deposition in monolayer cell cultures as well as in alginate-encapsulated bone spheroids. We observed that for encapsulated spheroids, colorimetric and fluorescent assays were not consistently reliable. We then explored the potential of SRS microscopy for label-free imaging of mineralized ECM. Our findings show that SRS microscopy can be successfully used to assess the mineralization state of the spheroid, by probing the vibrational energy transitions of phosphate groups (PO_4_^3–^), that are characteristic of calcium phosphate mineralization. In addition, SHG microscopy provides complementary information on the organization of the collagen matrix in the 3D construct. Together, SRS and SHG microscopy can be used to obtain complete information on both the mineralization state and organization of the ECM in the spheroid.

## 2 Results and discussion

### 2.1 Assessment of matrix mineralization in monolayer cell cultures and alginate-encapsulated spheroids via colorimetric and fluorescent assays

The deposition of mineralized matrix in monolayer cell cultures is typically assessed using staining dyes, such as Alizarin Red stain (ARS), calcein or commercial assays like OsteoImage™. However, the applicability of these methods can be limited in 3D cell cultures, due to the restricted dye penetration and signal attenuation in 3D samples [10, 41]. While bone spheroids are widely used in BTE research, microscopy-based visualization of cells and the mineralized ECM is challenging when not relying on techniques that require sample sectioning and chemical processing [12]. To evaluate if traditional colorimetric assays were suitable for detecting mineralization in 3D spheroid model, ARS, calcein and OsteoImage™ were applied to both monolayer cultures and alginate-encapsulated spheroids.

MC3T3-E1 subclone 4 cells cultured as monolayers in osteogenic media (OM) for 4 weeks exhibited extensive mineral deposition, which appeared as dark deposits when observed with bright-field microscopy (Figure 1A). These darker areas were positively stained with ARS, calcein and OsteoImage™ (Figure 1A), confirming the presence of mineralized matrix. In contrast, no mineralization was observed for cell cultures in regular media (RM). These results confirm that traditional staining techniques are reliable for the detection of matrix mineralization in 2D cell cultures. Moreover, mineral deposition could also be visualized without the employment of external dyes by simple bright-field microscopy. These same assays were applied to the alginate-encapsulated bone spheroids cultured for 4 weeks. Contrary to monolayer cell cultures, bright-field images did not provide information on the mineralization status of the samples (Figure 1B). ARS staining was positive in OM-cultured spheroids, whereas RM-cultured spheroids remained unstained (Figure 1B). In contrast to ARS, detectable fluorescence signal was observed in both RM and OM cultures supplemented with calcein, although a stronger signal was observed in OM-cultured samples (Figure 1B). In addition, some background signal originating from the alginate gel was observed, indicating non-specific staining of the alginate hydrogel. Although OsteoImage™ has been successfully used in previous studies to detect matrix mineralization in osteogenic spheroids [42], in our system, it did not provide reliable results. The alginate gel was stained with OsteoImage (Figure 1B), hindering the imaging of the encapsulated spheroids. These findings suggest that while traditional colorimetric and fluorescent assays are valid methods for mineral detection in monolayer cell cultures, they do not translate properly to hydrogel-embedded spheroids. Among these methods, ARS showed the most consistent results, in alignment with other studies where ARS was used to evaluate the mineralization state of the spheroid [43–46]. However, although ARS appears to be the most consistent method, its application in 3D cell models still suffers due to possible issues in dye penetration [26], highlighting the need for a label-free approach.

**Figure 1.**
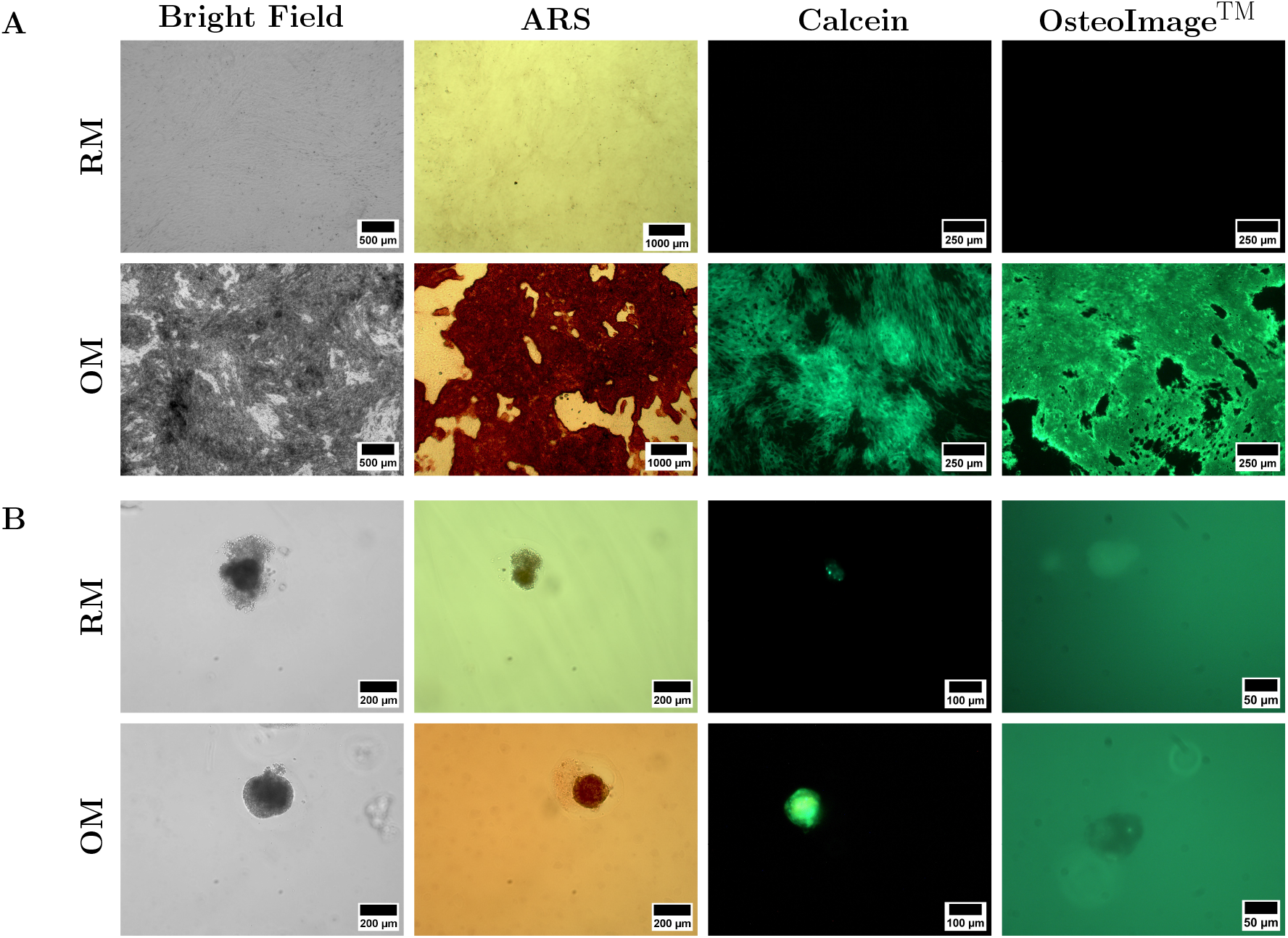
Analysis of mineralized matrix deposition in MC3T3-E1 monolayer cell cultures (A) and MC3T3-E1 alginate-encapsulated bone spheroids (B) after 4 weeks of culture, using colorimetric and fluorescent assays. In monolayer cell cultures (A), bright field images show the presence of dark material deposited on top of the cell layer in OM-cultured cells. These deposits were positively stained for ARS, calcein and OsteoImage™, confirming that matrix mineralization was taking place. RM-cultured cells were negative for all staining methods. In spheroids (B), bright field images did not provide any information regarding the mineralization state of the sample. While OM-cultured spheroid were positively stained for ARS, fluorescent signal was detected in both RM- and OM-cultured spheroids supplemented with calcein, with a stronger signal observed in OM-cultured spheroids. OsteoImage™ stained samples did not provide any relevant information.

### 2.2 Application of Coherent Raman Scattering microscopy for mineralization analysis

To address the need for label-free analysis of bone ECM in 3D, we employ Stimulated Raman Scattering (SRS) microscopy in combination with Second Harmonic Generation (SHG) microscopy and image mineralized ECM in alginate-encapsulated bone spheroids. To date, most SRS-based studies in cellular systems have focused on the high wavenumber region to primarily image lipids and proteins in single-cell studies or monolayer cell cultures [38, 47–50], and in spheroids to detect drug uptake [51]. However, the application of SRS to specifically probe calcium phosphate vibration in complex 3D cell models remains unexplored, with only limited publications addressing this in cellular models [52]. SRS microscopy offers several advantages. First, it requires minimal sample preparation, as imaging can be performed on both live and fixed samples (e.g., with paraformaldehyde). No sectioning is required, allowing to image full, intact construct. Because SRS and SHG are label-free, the signal is obtained exclusively from endogenous molecules and structures that are actually present in the sample, reducing the background and the risk of false-positive or false-negative results, as in the case of non-penetrating dyes. In addition, SRS allows for imaging of different molecular components by selecting characteristic Raman vibrations, without the need for multiple exogenous labels. Lastly, SRS and epi-SHG imaging can be performed simultaneously, allowing to visualize the two main ECM components at the same time.

#### 2.3.1 Detection of mineral deposits in monolayer cell cultures by SRS

To evaluate the ability of SRS microscopy to detect mineralized deposits, we first imaged MC3T3-E1 subclone 4 cells cultured as monolayers for 4 weeks in OM. The imaging was performed in transmission mode using a Raman shift of 960 cm^−1^, corresponding to the symmetric stretching vibration of the phosphate groups (PO_4_^3–^) [53]. Simultaneously, backscattered SHG (epi-SHG) microscopy was performed to assess collagen deposition.

SHG microscopy revealed the presence of a uniform collagen layer covering the cells (Figure 2B), with some areas where collagen looked more densely packed. SRS analysis on these areas revealed a positive SRS signal for the phosphate groups (Figure 2A), suggesting that the mineral deposition was taking place in these collagen-dense areas. To support these observation, transmission electron microscopy (TEM) performed on parallel cultures at a time point of 3 weeks revealed the presence of an organized, collagen-rich ECM, with electron-dense mineral deposits embedded within the ECM (Figure 3), confirming that MC3T3-E1 cells cultured in osteogenic conditions deposit mineral in a collagen-rich ECM. These results agree with the results from the colorimetric and fluorescent assays on 2D cultures.

**Figure 2.**
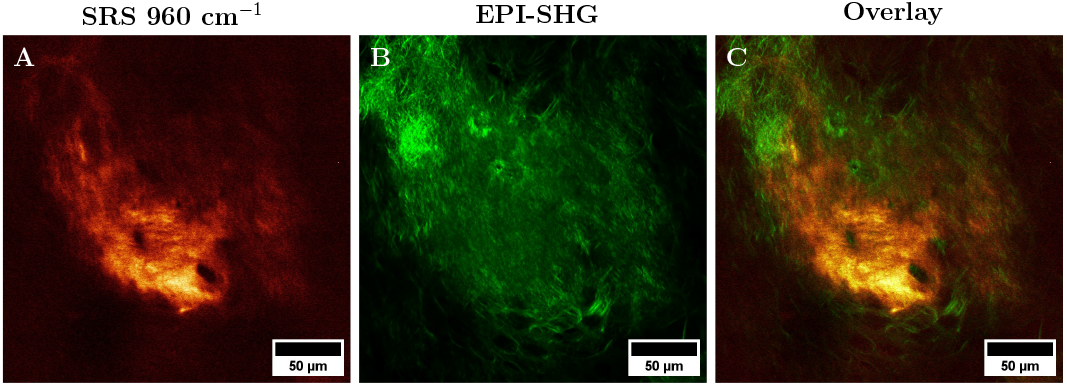
SRS and epi-SHG images of mineral deposits and collagenous matrix deposited by MC3T3-E1 cells after 4 weeks of culture in OM. A) SRS image at a Raman shift of 960 cm^−1^ to detect the presence of mineral deposits by probing the phosphate groups. B) Epi-SHG image of the deposited collagen matrix (green). C) Overlay of the SRS and epi-SHG images to visualize the localization of the mineralized region in relation to the localization of the collagen matrix.

**Figure 3.**
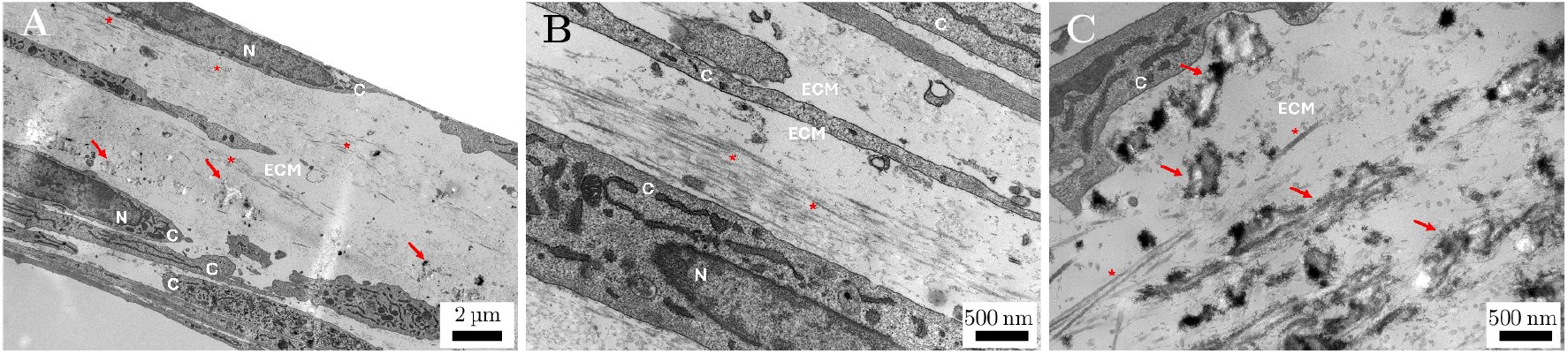
TEM images of mineralized ECM in MC3T3-E1 monolayer cell cultures after 3 weeks of culture in OM. A) Low magnification image showing the cell (C), nuclei (N), and ECM organization, with collagen fibrils (*) and mineral deposits (arrow). B) High magnification image of an area of highly organized collagen fibrils (*), cells (C) and nuclei (N). C) High magnification image showing electron-dense mineral deposits (arrow) within the deposited collagen (*), indicating mineralization of the ECM.

Initial attempts to image monolayer cells cultured on polystyrene imaging slides resulted in a weak SRS signal, while epi-SHG produced satisfactory signals. Transferring the same cell layer onto glass cover-slips substantially improved the signal quality, resulting in an optimal signal being obtained both in the SRS and epi-SHG channels. This difference between polystyrene and glass can be attributed to the optical properties of the polystyrene slide and the detection of the SRS signal in the transmitted light direction. In addition, although polystyrene exhibits the main Raman peak at 1001 cm^−1^ [54], background contributions from the substrate could still be observed, thus influencing the observed SRS signal. This observation highlights the importance of proper substrate selection prior to imaging.

#### 2.2.2 Combination of SRS and epi-SHG imaging for the analysis of mineralized ECM in alginate-encapsulated bone spheroids

The same approach was applied to analyze the ECM organization and mineralization in spheroids. The SRS image at 960 cm^−1^ of OM-cultured spheroids revealed the presence of brighter regions, which correspond to phosphate-rich areas (Figure 4A). To confirm that the signal in these regions was specific for CaP, Raman spectra in the range between 900 to 1000 cm^−1^ were acquired. Six regions-of-interests (ROI) were selected based on the SRS signal intensity, including regions with strong and weak signal, as well as from the surrounding alginate hydrogel. The normalized Raman spectra revealed the presence of a distinct peak at 960 cm^−1^ in regions with SRS signal, while no such peak was observed in regions that did not exhibit a SRS signal or in the surrounding hydrogel (Figure 4B). It is important to notice that the Raman spectra obtained using the SRS are equivalent to spontaneous Raman scattering spectra[35, 55]. The possibility of obtaining a Raman spectra allows for an additional chemical validation of the SRS images. The same spectral analysis was performed on RM-cultured spheroids, and confirmed the absence of phosphate-rich deposits, further supporting the specificity of the SRS detection of the mineralized deposits (Figure S1 A and B).

**Figure 4.**
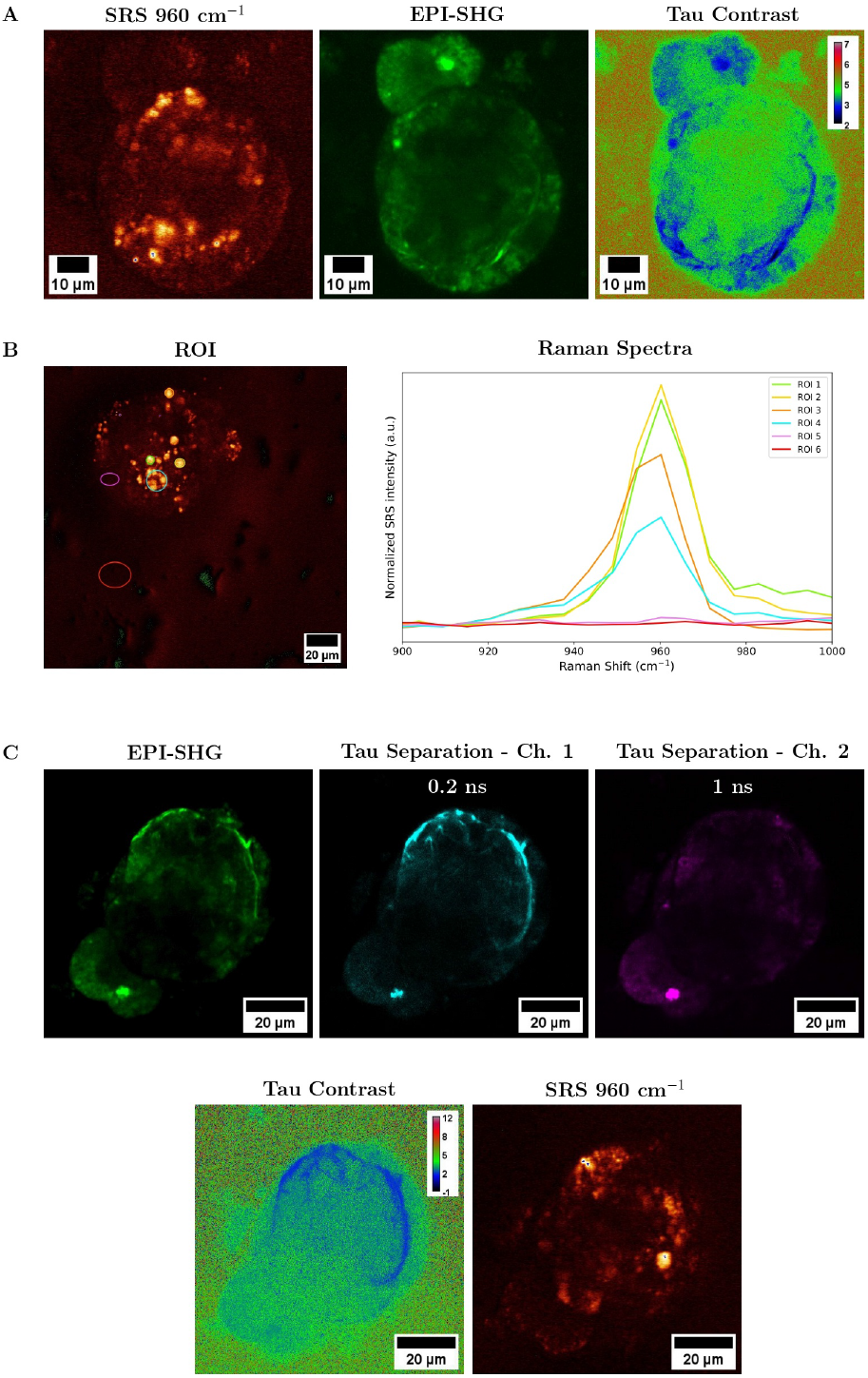
SRS and SHG analysis of mineralized ECM in alginate-encapsulated bone spheroids cultured for 4 weeks in OM. A) Combined SRS and epi-SHG images of a representative spheroid. The SRS image was obtained at Raman shift of 960 cm^−1^ to detect phosphate groups associated with mineralized deposits. The epi-SHG signal consist of contributions from the collagenous matrix and cellular auto-fluorescence; the signals were distinguished using the TauContrast function. The calibration bar represents the average arrival-time in nanoseconds. B) Raman spectra acquired in the 900 to 1000 cm^−1^ range and corresponding SRS image, showing the selected regions of interest (ROIs). C) Epi-SHG images showing the signal contributions and the corresponding Tau-Separated channels, allowing the discrimination of collagen-derived SHG and auto-fluorescence based signal based on effective photon lifetime. The TauContrast image and SRS image at 960 cm^−1^ were collected simultaneously. The calibration bar in the TauContrast image represents the average arrival-time in nanoseconds.

The mineral deposits in the spheroids were mostly co-localized with the collagen matrix, as revealed by epi-SHG microscopy. These results were also supported by TEM imaging performed on parallel spheroids samples. The images show a well-organized spheroid, composed of cells embedded in a dense collagenous ECM with organized fibrillar structure (Figure 5 A, B, C). Small mineral deposits are visible in the ECM (Figure 5 D). The morphology of these deposits is consistent with results observed from previous reports [6, 56, 57]. The limited amount of mineral detected using TEM is most likely caused by the sample preparation, which can lead to partial loss or dissolution of CaP[57]. This aspect highlights the potential and the advantage of SRS for studies on early stages of mineral deposition, as it allows the detection of phosphate-rich areas in intact 3D constructs. Although epi-SHG is well suited for the detection of the collagen matrix in alginate-encapsulated bone spheroids as we demonstrated in our previous publication [40], in the SHG images collected in this work (Figure 4A), auto-fluorescence originating from cells was also detected in the epi-SHG channel. This was due to the broad detection bandwidth of the detection filter (465/170 nm). To investigate whether these signals could be distinguished based on their temporal characteristics, the TauContrast and TauSeparation functionality of the Leica Stellaris 8 microscope were used to achieve photon arrival-time-based discrimination between the two signals. TauContrast measures the average photon arrival-time, while Tau Separation divides the photons into two channels based on their effective arrival-time, under the assumption that in the same pixel different populations of photons coexist. Fast processes, such as SHG will have a very short arrival-time (and corresponding short apparent lifetime), whereas the fluorescence signal should have a longer arrival-time due to the lifetime of the excited state of the fluorescent molecules. Considering the physical difference of the two processes observed in the SHG channel, TauContrast and TauSeparation can help separate the SHG signal from the auto-fluorescence. In this context, it is important to consider that in TauContrast, both fast SHG and slow auto-fluorescence photons are averaged together, meaning that the slow photons could influence the apparent lifetime. For this reason, TauContrast was first used to obtain a general overview of the lifetime differences in the SHG channel, showing the regions with collagen and those characterized by auto-fluorescence. Subsequently, TauSeparation was applied to divide the signals into two different channels (Figure 4 C), allowing for a precise and effective measurement of the photon lifetime. An effective separation of the collagen and auto-fluorescence signal was obtained by TauSeparation, where collagen structures were detected in the short arrival-time channel, which revealed an effective photon arrival time of 0.2 ns, while the auto-fluorescence was limited to the long arrival-time channel, with an effective arrival time of 1 ns (Figure 4C).

**Figure 5.**
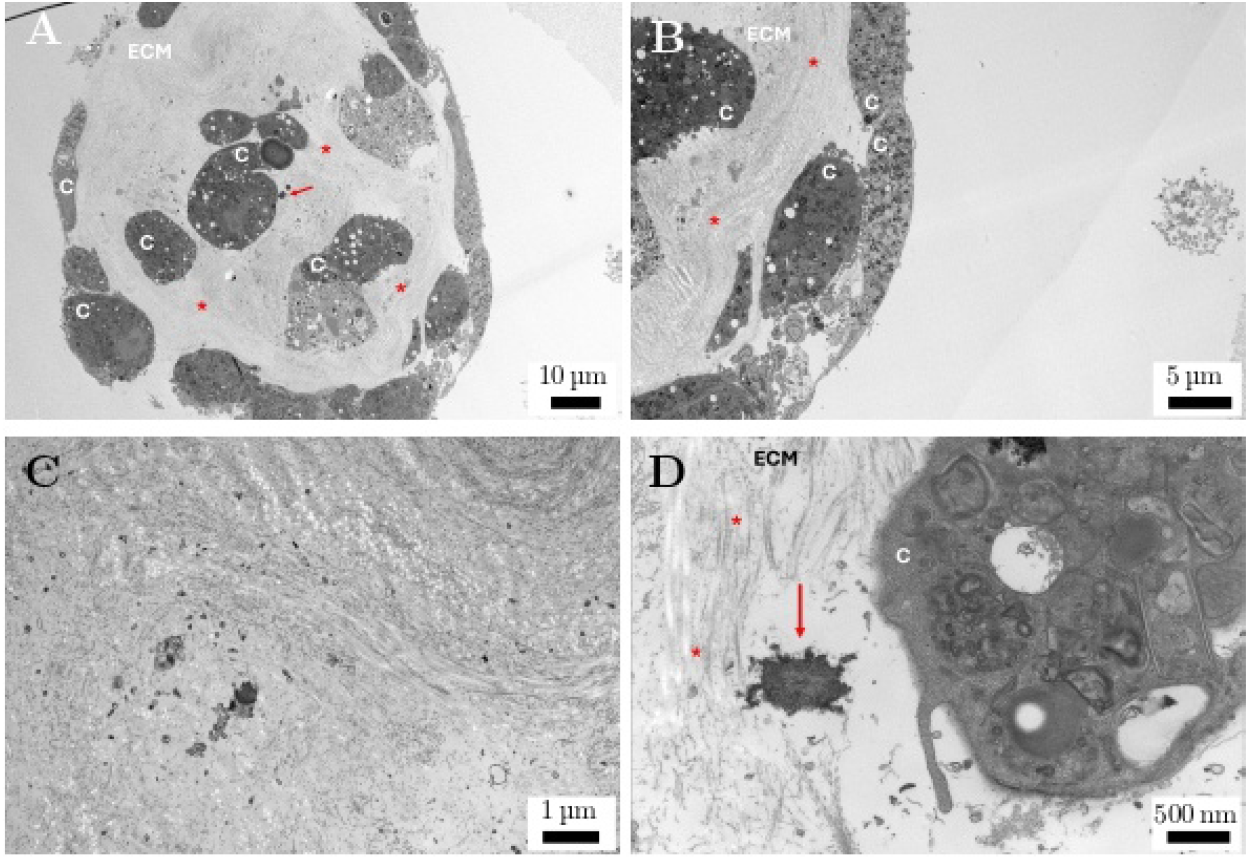
TEM images of mineralized ECM in alginate-encapsulated bone spheroids after 6 weeks of culture in OM. A) Low magnification image showing the general spheroid organization, with the cells (C), extracellular matrix (ECM), collagen (*) and small mineral deposits (arrow). B) High magnification image of an area of highly organized collagen matrix (*) and cells (C). It can be observed the presence of a small vesicle in the pocket area between the spheroids and the alginate hydrogel. For more info and details, refer to [40]. C) High magnification image showing an area of ECM rich in organized collagen. D) High magnification image of an area in the spheroid showing the presence of a cell (C), organized collagen (*) and an electron-dense mineralized deposit (arrow), indicating mineralization of the ECM.

To further confirm that the observed signal was specific to the mineralized deposits, the same analysis was performed on RM-cultured spheroids. Only long lifetime values were detected in the SHG TauContrast and TauSeparation channels, and no SRS signal at 960 cm^−1^ was observed (Figure S1). These results confirm that the SRS signal observed in OM-cultured spheroid arises from mineralized deposits. Epi-SHG imaging confirmed the absence of collagen deposition (Figure S1). These observations confirmed that while TauContrast is a useful tool to quickly assess the distribution of fast and slow components, TauSeparation provides a more precise measurement.

#### 2.2.3 Validation of label-free SRS imaging through comparison with calcein fluorescence

To evaluate the specificity of SRS to detect mineralized deposits, SRS was performed simultaneously with two-photon fluorescence (TPF) on calcein-stained spheroids, using the epi-SHG detection channel. The TPF was collected using the same epi-detection pathway (465/170 nm), resulting in the simultaneous detection of both SHG and TPF signal. The distribution of the SRS phosphate signal was then compared to the calcein signal by overlaying the two channels to assess co-localization of the mineralized regions identified by the two different imaging methods and to evaluate if the phosphate regions detected via SRS were consistent with the areas that were labeled using calcein.

In the SRS images of calcein-supplemented OM-cultured spheroids, phosphate-rich regions appear as bright areas, as observed during the previous analysis (Figure 6A). In the corresponding SHG/TPF image, multiple processes contributed to the observed signal. The collagen fibers were imaged via SHG (as an SHG detector was used to perform TPF imaging) and could be identified by their typical fibrous structure. The other signal observed was a fluorescence signal that comprised both calcein fluorescence and the auto-fluorescence, as discussed above. In the outer layer of the spheroid, region showing calcein fluorescence spatially overlapped with the SRS-positive regions (Figure 6 A, B and C), indicating that these SRS-positive areas correspond to calcein-stained mineralized deposits. In contrast, in the spheroid core region, a strong SRS signal was detected, but no corresponding calcein fluorescence was observed, suggesting the presence of mineralized deposits that were not been labeled by calcein. The TauContrast image (Figure S2) confirms the co-localization between SRS-positive regions and regions with longer average photon arrival-time, originating from calcein fluorescence. The non-overlapping fluorescent signal was attributed to auto-fluorescence. To further confirm that the fluorescent signal of calcein-stained deposits was originating from the dye and was not caused by auto-fluorescence, unstained OM-cultured samples were imaged and compared. Similarly to the calcein-stained sample, the SHG/TPF image is characterized by the presence of the deposited collagen fibrils and auto-fluorescence (Figure 6E). However, no overlap was observed between the auto-fluorescent regions and the SRS-positive regions. These results confirm that the overlapping regions observed in the calcein-stained OM-cultured spheroid corresponded to calcein-stained mineral deposits. Calcein-stained RM-cultured spheroids did not show any SRS-positive region (Figure 6G). Furthermore, the TPF image was characterized by auto-fluorescence and by the absence of collagenous matrix (Figure 6H). While the observed overlap confirms that both techniques allow the identification of mineralized deposits, it is important to notice that calcein labels any calcium source, making it non-specific for mineralized deposits. In contrast, SRS directly probes the specific vibrations of the phosphate groups in CaP deposits, making it a highly-specific, label-free method for mineralization analysis. In addition, SRS enables imaging of the full spheroid volume, including the core region, that showed no calcein fluorescence despite being SRS-positive, making it a more robust and reliable approach for assessing not only mineral deposition but also to perform a comprehensive characterization of the 3D aggregate.

**Figure 6.**
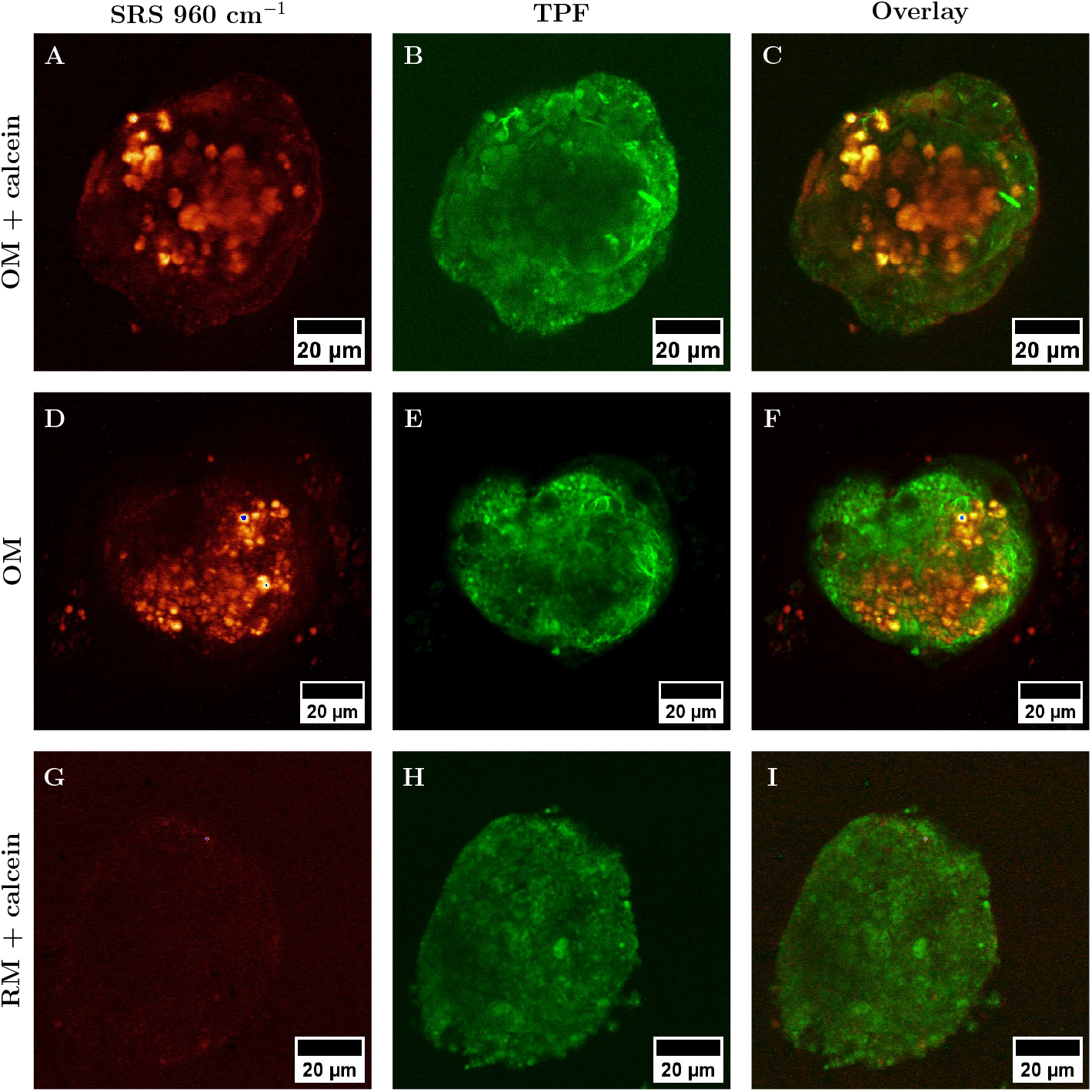
Validation of label-free SRS imaging through comparison with calcein fluorescence. Combined SRS, two-photon fluorescence and overlay images. A-C) Representative bone spheroid cultured in calcein-supplemented OM for 4 weeks. D-F) Representative bone spheroid cultured in OM for 4 weeks. G-I) Representative spheroid cultured in calcein-supplemented RM for 4 weeks. A-D-G) SRS images were acquired at a Raman shift of 960 cm^−1^ to detect the phosphate groups associated with mineralized deposits. B-E-H) Two-photon fluorescent images containing contributions from the collagen matrix (B-E), auto-fluorescence (B-E-H) and calcein (B). C-F-I) Overlay of the two channel for analysis of possible overlapping regions between calcein-positive areas and SRS-positive areas. All images were acquired under identical conditions.

#### 2.2.4 Comparison of SRS and CARS for localization of lipid-rich regions for cells localization

Alginate-encapsulated bone spheroids were simultaneously imaged using epi-CARS and SRS, targeting the C-H vibration region in the CH_2_ stretching (2857 cm^−1^), which was used to provide a general overview of the cell distribution in the 3D structure by visualizing lipid-rich regions. Comparison between the two channels showed that while many regions overlapped, in the CARS channel there was an additional signal that was not observed in the SRS channel (Figure 7A and B). To further confirm that the nature of this signal was not CARS-related and therefore not chemically specific to the lipids, a second image was acquired with the Stoke laser turned off. In this case, no signal was detected in the SRS channel (data not shown), while in the CARS channel only the additional signal caused by two-photon fluorescence was present (Figure 7C), confirming its non-CARS related nature. Thus, SRS imaging resulted in a more reliable localization of regions of interest without non-resonant background when compared to CARS.

**Figure 7.**
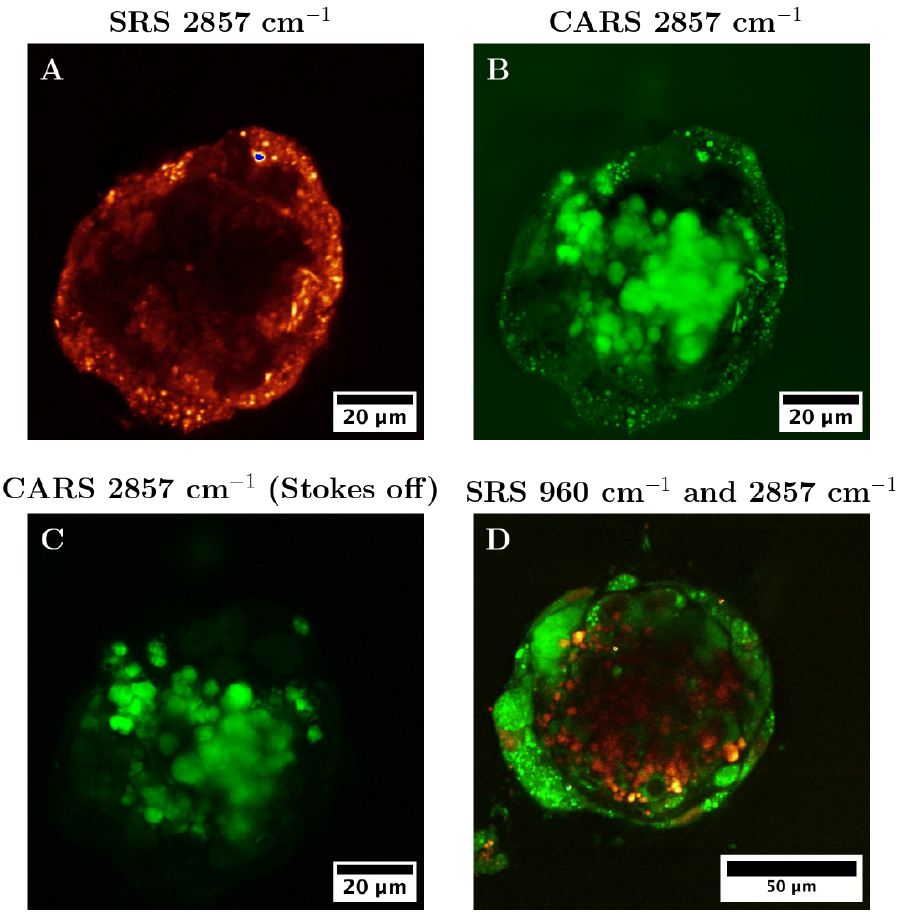
Comparison between SRS and CARS imaging of lipids-rich regions. A) SRS image at a Raman shift of 2857 cm^−1^. B) CARS image at a Raman shift of 2857 cm^−1^. C) CARS image at a Raman shift of 2857 cm^−1^ with the Stokes laser turned off. D) SRS image of a representative bone spheroid. Images were acquired sequentially at 960 cm^−1^ (red) to visualize mineralized regions and 2857 cm^−1^ (green) to detect lipid-rich areas, highlighting the spatial distribution of these two components within the spheroid.

Simultaneous SRS imaging at 960 cm^−1^ and 2857 cm^−1^ was used to successfully obtain an overview of the localization of the cells in the spheroids, using lipids-rich regions, and of the mineralized deposits (Figure 7D), highlighting the potential of SRS for extensive spheroid characterization, provided that appropriate Raman shifts are selected for the structure or molecules of interest.

## 3 Conclusion

In this study, we demonstrated that SRS microscopy is a promising approach to perform label-free imaging of matrix mineralization in alginate-encapsulated bone spheroids. Traditional colorimetric and fluorescent assays, such as ARS and calcein, were first applied on monolayer cell cultures to assess mineral deposition. After confirming that these methods were not well-suited for investigating matrix mineralization in 3D cell spheroids, SRS was employed to perform a label-free imaging of the mineralized deposits. SHG was further used to perform label-free imaging of the collagenous matrix. Our results demonstrate that together, SRS and SHG allow for high-resolution, label-free and chemically specific characterization of both matrix mineralization and organization in the spheroids, highlighting also the potential of SRS for a comprehensive, chemically-specific and non-invasive characterization of the 3D cell aggregate.

## 4 Material and methods

### 4.1 Cell culture

MC3T3-E1 subclone 4 (ATCC, CRL-2593) were cultured in standard conditions at 37 ^°^C and 5% CO_2_ in minimum essential alpha medium (MEM-*α* without ascorbic acid; ThermoFisher, A1049001) supplemented with 10% fetal bovine serum (FBS; Sigma-Aldrich). To induce osteogenic differentiation, the growth medium was supplemented with 50 µg*/*mL of L-ascorbic acid 2-phosphate sesquimagnesium salt hydrate (Sigma-Aldrich, A8960) and 5 mM of *β*-glycerophosphate disodium salt pentahydrate (Sigma-Aldrich).

For colorimetric and fluorescent assays, MC3T3-E1 were cultured in 6 well-plates. One well was maintained unstained for SRS and CARS microscopy. For SRS and CARS microscopy, the cell layer was moved from the well in between two glass coverslips to ensure optimal imaging. Bright-field images of monolayer cells were obtained using the Nikon ECLIPSE TS100 microscope and a 4X objective lens (Nikon, Tokyo, Japan).

### 4.2 Spheroids formation

Spheroids were obtained using the micro-mold technique [58] as described before[40]. Briefly, 1.5% agarose (Sigma-Aldrich) molds were made from #24-96 silicon molds (MicroTissues 3D Petri Dish micro-mold spheroids) and were placed in a 24-well plate. The cells were collected and resuspended in 1 mL of RM, and 75 µL of cell suspension were added to each mold, followed by 1 mL of RM added into the well. After overnight incubation, the molds were placed upside down in a new 24-well plate and centrifuged to allow spheroid collection. The molds were removed and the media with spheroids was collected in a 15 mL tube and centrifuged. The sedimented spheroids were retrieved and resuspended in a solution made of an equal volume of RM and sterile filtered 2% Alginate (Alg) solution (G fraction: 0.68, GG fraction: 0.57, Molecular weight: 250 000 g*/*mol, [Novamatrix (Sandvika, Norway)]). 150 µL of the spheroids solution was added into glassed-bottomed Petri Dishes (35 mm Dish with 14 mm bottom well, #1.5 glass – 0.16 - 0.19, Cellvis, Cat. #D35-10-1.5-N), covered with a GN-6 Metricel® 0.45 µm-47 mm sterile membrane and gelation was induced by adding 50 mM CaCl_2_ on top of the membrane for 5 minutes. After gelation, both the CaCl_2_ solution and the membrane were removed. The alginate disks containing spheroids were kept in 2 mL of media. Media was changed every second or third day. Spheroids were maintained with the same standard conditions as the monolayer cell cultures. Osteogenic media (OM) was used to induce osteogenic differentiation of the spheroids in the alginate disks. Bright-field images of spheroids were obtained using the Motic AE31E microscope and a 20X/0.3 objective lens.

### 4.3 Assessment of CaP deposition using colorimetric assays and fluorescent dyes

#### 4.3.1 Alizarin Red Staining

Alizarin Red Stain (ARS) was used to stain CaP deposits in monolayer cell cultures and spheroids. Monolayer cells cultured for 4 weeks in RM and OM were first rinsed with PBS and then fixed using 4% PFA for 20 minutes. After rinsing with PBS three times for 5 minutes, the samples were incubated with 2% w/v ARS solution (Sigma-Aldrich, A5533) for 20 minutes, covered from light. The samples were then washed with MilliQ water to remove the excess staining solution and imaged.

Alginate-embedded spheroids cultured for 4 weeks in RM and OM were first washed with 5 mM BaCl for 5 minutes. The samples were then fixed using 4% PFA for 20 minutes, and rinsed three times with PBS for 5 minutes. The alginate gels were incubated with 1 mL of 2% w/v ARS solution for 20 minutes, protected from light. The hydrogels were then washed with MilliQ water to remove excess staining solution and imaged. To speed up the washing process, the gels were placed on a shaker and kept there until the complete release of the excess dye.

Both monolayer cell cultures and spheroids were imaged using the Motic AE31E microscope and a 20X/0.3 objective lens.

#### 4.3.2 Calcein staining

On the sample preparation day, one well containing 2D cell cultures and one alginate disk with embedded spheroids were randomly selected for staining with Calcein (ex/em 488/520; Invitrogen™). Samples from both RM and OM culture conditions were included. Calcein was added to the RM and OM media at a final concentration of 1 µg*/*mL, and the samples were cultured in the calcein-supplemented media for the designated incubation period. On imaging day, both 2D cell cultures and alginate-encapsulated spheroids were fixed as mentioned in the ARS section. After washing, the samples were imaged using the Nikon ECLIPSE TS100 microscope, equipped with a 10X/0.3 objective lens for monolayers and 20X/0.45 for spheroids, and a FITC filter (ex/em 490/520 nm).

#### 4.3.3 Osteoimage™ staining

OsteoImage™ Mineralization Assay (Lonza) was used to stain the CaP deposits, as per manufacturer’s instruction. Prior to staining, monolayer cell cultures and alginate-encapsulated spheroid were fixed as described in the ARS section. Samples were then imaged using the Nikon ECLIPSE TS100 microscope, equipped with a FITC filter (ex/em 490/520 nm).

### 4.4 Coherent Raman Scattering

Coherent Raman Scattering (CRS) microscopy as well as multiphoton fluorescence/second harmonic generation (SHG) microscopy were carried out on an inverted Leica Stellaris 8 CRS microscope (Leica Microsystems), equipped with a picoEmerald S dual beam infrared laser (APE), providing two pulsed and synchronized beams, a fixed Stokes beam (1032 nm) and a tunable pump beam (720 nm – 980 nm). Stimulated Raman Scattering (SRS) was recorded in the forward or transmission direction as energy transfer between the modulated Stokes beam and the pump beam by a photodiode. Coherent Anti-Stokes Raman Scattering (CARS) and SHG signals were detected in the reflected (epi) direction via photon-counting detectors (HyD, Leica Microsystems). Emission bandpass filters of 465/170 nm and 670/125 nm were used for SHG and CARS detection, respectively. Images were acquired using a 25x water immersion objective (HC FLUOTAR L 25x/0.95 W VISIR, Leica Microsystems) and in the case of the transmitted light direction, a high numerical aperture condenser (1.40 OIL S1, used with water immersion, Leica Microsystems). Samples (both monolayer cell cultures and alginate-embedded spheroids) were imaged between two glass coverslips, and the condenser was aligned for Köhler illumination. Imaging parameters were chosen as following. SRS measurements were collected using a pump beam at 938.7 nm (the energy difference to the Stokes beam corresponding to Raman shift 960 cm^−1^) for detection of phosphate groups, and at 796.8 nm (2857 cm^−1^) for lipids. For spectral scans (xyΛ), the step size was fixed to 0.5 nm and the field of view was chosen to include all relevant areas, such as spheroid, pocket and alginate. The collected Raman spectra were normalized using the Local Reference Normalization in the 900-920 cm^−1^ region. Both types of samples were prepared and fixed as described in the ARS section.

### 4.5 Transmission Electron Microscopy

For mineral deposition analysis, TEM was performed on MC3T3-E1 cells cultured either as monolayer and as embedded spheroids. Monolayer samples were cultured on Aclar film for 3 weeks in OM, while spheroids were maintained in standard conditions in OM for 6 weeks. The same sample preparation process was used for both types of samples. After 3 weeks for monolayer cell cultures and 6 weeks for spheroids, the samples were fixed in 2% formaldehyde, 2.5% glutaraldehyde, and 0.025% CaCl_2_ in 0.1 M cacodylate buffer (pH = 7.4) overnight at room temperature (RT). The samples were washed with 0.1 M cacodylate buffer with 0.025% CaCl_2_ twice for 15 minutes. At this stage, the alginate disk was cut into smaller pieces to facilitate chemical penetration during subsequent sample preparation. Samples were then post-fixed in 2% osmium tetroxide, 0.025% CaCl_2_ and 1.5% potassium ferrocyanide for 45 minutes. The samples were washed again with the same washing solution as mentioned before and dehydrated in increasing ethanol series (50%, 70%, 90%) for minutes each. The samples were followed by uranyl acetate (UA) staining (4% UA in 50% ethanol) for 1 hour and 30 minutes. Lastly, the samples were washed first in absolute ethanol and then in acetone and embedded in a pure acetone and epoxy (AGAR) resin mix in a ratio of 2:1, 1:1 and 1:2 for 45 minutes each. The samples were then incubated in epoxy resin overnight. The samples were embedded in fresh resin and polymerized at 60 ^°^C overnight. At this point, the Aclar was removed from the monolayer cells samples. The samples (both monolayer and spheroids) were left to further polymerize at 60 ^°^C for further 24 hours. The samples were sectioned with a Leica UC7 Ultramicrotome at 60 nm. The sections were collected on copper grids (Gilder) covered with a thin formvar-film. The sections were post-stained with 4% UA in 50% ethanol and 1% lead citrate in 0.1 M NaOH for 25 and 5 minutes respectively. Monolayer and spheroid sections were imaged using a FEI Tecnai 12 with a tungsten filament, with an acceleration voltage of 80 kV.

## Acknowledgements

The authors thank BNMI for the financial support. We acknowledge Nan Tostrup Skogaker for training, technical assistance and access to the electron microscopy core facility, NTNU. We also acknowledge Stiftelsen Biopolymer for financial support. SRS, SHG and multiphoton fluorescence data were collected at the Center for Advanced Bioimaging (CAB) Denmark, University of Copenhagen, which is operated with funding from the Novo Nordisk Foundation (NNF23OC0082200).

## Supporting information

Supporting information: Figure S1: SRS Analysis of mineralized ECM in alginate-encapsulated bone spheroids cultured for 4 weeks in RM; Figure S2: Validation of label-free SRS imaging through comparison with calcein fluorescence. Combined SRS, two-photon fluorescence and Tau Contrast images.

## Notes

### Competing Interest Statement

The authors have declared no competing interest.

## References

(1) Caetano-Lopes, J.; Canhão, H.; Fonseca, J. E. Acta Reumatol. Port. 2007, 32, 103–110.

(2) Oliveira, C. S.; Leeuwenburgh, S.; Mano, J. F. APL Bioengineering 2021, 5, 041507.

(3) Mansour, A.; Mezour, M. A.; Badran, Z.; Tamimi, F. Tissue Engineering Part A 2017, 23, 1436–1451.

(4) Clarke, B. Clinical Journal of the American Society of Nephrology 2008, 3, S131–S139.

(5) Brochado, A. C. B.; Silva, D. C.; Silva, J. C. d.; Lowenstein, A.; Gameiro, V. S.; Mavropoulos, E.; Mourão, C. F.; Alves, G. G. Applied Sciences 2023, 13, 1602.

(6) Addison, W.; Nelea, V.; Chicatun, F.; Chien, Y.-C.; Tran-Khanh, N.; Buschmann, M.; Nazhat, S.; Kaartinen, M.; Vali, H.; Tecklenburg, M.; Franceschi, R.; McKee, M. Bone 2015, 71, 244–256.

(7) Mertz, E. L.; Makareeva, E.; Mirigian, L. S.; Leikin, S. JBMR Plus 2023, 7, e10701.

(8) Yoon, H.; Park, S. G.; Shin, H.-R.; Kim, K.-T.; Cho, Y.-D.; Moon, J.-I.; Kim, W.-J.; Ryoo, H.-M. Bone 2025, 194, 117442.

(9) Yuste, I.; Luciano, F.; González-Burgos, E.; Lalatsa, A.; Serrano, D. Pharmacological Research 2021, 169, 105626.

(10) Urzì, O.; Gasparro, R.; Costanzo, E.; De Luca, A.; Giavaresi, G.; Fontana, S.; Alessandro, R. International Journal of Molecular Sciences 2023, 24, 12046.

(11) Griffith, L. G.; Swartz, M. A. Nature Reviews Molecular Cell Biology 2006, 7, 211–224.

(12) Yun, C.; Kim, S. H.; Kim, K. M.; Yang, M. H.; Byun, M. R.; Kim, J.-H.; Kwon, D.; Pham, H. T. M.; Kim, H.-S.; Kim, J.-H.; Jung, Y.-S. International Journal of Molecular Sciences 2024, 25, 2512.

(13) Gionet-Gonzales, M. A.; Leach, J. K. Biomedical Materials 2018, 13, 034109.

(14) Nitschke, B. M.; Beltran, F. O.; Hahn, M. S.; Grunlan, M. A. Journal of Materials Chemistry B 2024, 12, 2720–2736.

(15) White, K.; Chalaby, R.; Lowe, G.; Berlin, J.; Glackin, C.; Olabisi, R. Polymers 2021, 13, 2274.

(16) Verdugo-Avello, F.; Wychowaniec, J. K.; Villacis-Aguirre, C. A.; D’Este, M.; Toledo, J. R. Lab on a Chip 2025, 25, 806–836.

(17) Mao, W.; Bui, H.-T. D.; Cho, W.; Yoo, H. S. Advanced Drug Delivery Reviews 2023, 201, 115074.

(18) Krafft, C. Journal of Biomedical Optics 2012, 17, 040801.

(19) Ember, K. J. I.; Hoeve, M. A.; McAughtrie, S. L.; Bergholt, M. S.; Dwyer, B. J.; Stevens, M. M.; Faulds, K.; Forbes, S. J.; Campbell, C. J. npj Regenerative Medicine 2017, 2, 12.

(20) Charwat, V.; Schütze, K.; Holnthoner, W.; Lavrentieva, A.; Gangnus, R.; Hofbauer, P.; Hoffmann, C.; Angres, B.; Kasper, C. Journal of Biotechnology 2015, 205, 70–81.

(21) Ghita, A.; Pascut, F. C.; Sottile, V.; Notingher, I. The Analyst 2014, 139, 55–58.

(22) McManus, L. L.; Burke, G. A.; McCafferty, M. M.; O’Hare, P.; Modreanu, M.; Boyd, A. R.; Meenan, B. J. The Analyst 2011, 136, 2471.

(23) Gao, Y.; Xu, C.; Wang, L. RSC Advances 2016, 6, 61771–61776.

(24) Tarnowski, C. P.; Ignelzi, M. A.; Morris, M. D. Journal of Bone and Mineral Research 2002, 17, 1118–1126.

(25) Timlin, J. A.; Carden, A.; Morris, M. D.; Rajachar, R. M.; Kohn, D. H. Analytical Chemistry 2000, 72, 2229–2236.

(26) Kim, H.; Han, Y.; Suhito, I. R.; Choi, Y.; Kwon, M.; Son, H.; Kim, H.-R.; Kim, T.-H. Analytical Chemistry 2021, 93, 9995–10004.

(27) Liao, H.-X.; Bando, K.; Li, M.; Fujita, K. Analytical Chemistry 2023, 95, 14616–14623.

(28) Jamieson, L. E.; Harrison, D. J.; Campbell, C. J. Journal of Biophotonics 2019, 12, e201800201.

(29) Pettinato, G. et al. Science Advances 2021, 7, eabj2800.

(30) Bispo, D. S.; Graça, I. C. R.; Rodrigues, J. A.; Martins, J. T. S.; Nolasco, M. M.; Marques, M. P. M.; Nogueira, H. I. S.; Mano, J. F.; Oliveira, M. B.; Ribeiro-Claro, P. J. A.; Gil, A. M. Stem Cell Reviews and Reports 2025, DOI: 10.1007/s12015-025-10943-3.

(31) Brackmann, C.; Esguerra, M.; Olausson, D.; Delbro, D.; Krettek, A.; Gatenholm, P.; Enejder, A. Journal of Biomedical Optics 2011, 16, 021115.

(32) Cheng, J.-X.; Xie, X. S. The Journal of Physical Chemistry B 2004, 108, 827–840.

(33) Li, Y.; Shen, B.; Li, S.; Zhao, Y.; Qu, J.; Liu, L. Advanced Biology 2021, 5, 2000184.

(34) Müller, M.; Zumbusch, A. ChemPhysChem 2007, 8, 2156–2170.

(35) Brzozowski, K.; Pieczara, A.; Nowakowska, A. M.; Korona, W.; Orzechowska, B.; Firlej, J.; Wislocka-Orlowska, A.; Baranska, M. Optica 2025, 12, 11.

(36) Li, S.; Li, Y.; Yi, R.; Liu, L.; Qu, J. Frontiers in Physics 2020, 8, 598420.

(37) Cheng, Q.; Miao, Y.; Wild, J.; Min, W.; Yang, Y. Matter 2021, 4, 1460–1483.

(38) Freudiger, C. W.; Min, W.; Saar, B. G.; Lu, S.; Holtom, G. R.; He, C.; Tsai, J. C.; Kang, J. X.; Xie, X. S. Science 2008, 322, 1857–1861.

(39) Tipping, W. J.; Lee, M.; Serrels, A.; Brunton, V. G.; Hulme, A. N. Chemical Society Reviews 2016, 45, 2075–2089.

(40) Boscaro, D.; Wahlum, L. S.; Ullevålseter, M. E.; Strand, B. L.; Sikorski, P. Materials 2025, 18, 3538.

(41) Weiswald, L.-B.; Guinebretière, J.-M.; Richon, S.; Bellet, D.; Saubaméa, B.; Dangles-Marie, V. BMC Cancer 2010, 10, 106.

(42) Vermeulen, S.; Knoops, K.; Duimel, H.; Parvizifard, M.; Van Beurden, D.; López-Iglesias, C.; Giselbrecht, S.; Truckenmüller, R.; Habibović, P.; Tahmasebi Birgani, Z. Materials Today Bio 2023, 23, 100844.

(43) Moritani, Y.; Usui, M.; Sano, K.; Nakazawa, K.; Hanatani, T.; Nakatomi, M.; Iwata, T.; Sato, T.; Ariyoshi, W.; Nishihara, T.; Nakashima, K. Journal of Periodontal Research 2018, 53, 870–882.

(44) De Souza Castro, G.; De Souza, W.; Lima, T. S. M.; Bonfim, D. C.; Werckmann, J.; Archanjo, B. S.; Granjeiro, J. M.; Ribeiro, A. R.; Gemini-Piperni, S. Nanomaterials 2023, 13, 425.

(45) Xu, C.; Li, Z.; Kang, M.; Chen, Y.; Sheng, R.; Aghaloo, T.; Lee, M. Biomaterials 2025, 317, 123088.

(46) Sanchez, A. A.; Teixeira, F. C.; Casademunt, P.; Beeren, I.; Moroni, L.; Mota, C. Biofabrication 2025, 17, 025013.

(47) Yue, S.; Cheng, J.-X. Current Opinion in Chemical Biology 2016, 33, 46–57.

(48) Huang, J.; Zhang, L.; Shao, N.; Zhang, Y.; Xu, Y.; Zhou, Y.; Zhang, D.; Zhang, J.; Lee, H. J. Chemical & Biomedical Imaging 2025, 3, 15–24.

(49) Hill, A. H.; Fu, D. Analytical Chemistry 2019, 91, 9333–9342.

(50) Cheng, J.-X.; Yuan, Y.; Ni, H.; Ao, J.; Xia, Q.; Bolarinho, R.; Ge, X. Nature Methods 2025, 22, 912–927.

(51) Xu, F. X.; Sun, R.; Owens, R.; Hu, K.; Fu, D. Analytical Chemistry 2024, 96, 14480–14489.

(52) Mansfield, J.; Moger, J.; Green, E.; Moger, C.; Winlove, C. P. Journal of Biophotonics 2013, 6, 803–814.

(53) Movasaghi, Z.; Rehman, S.; Rehman, I. U. Applied Spectroscopy Reviews 2007, 42, 493–541.

(54) Mayorga, C.; Athalye, S. M.; Boodaghidizaji, M.; Sarathy, N.; Hosseini, M.; Ardekani, A.; Verma, M. S. Limit of detection of Raman spectroscopy using polystyrene particles from 25 to 1000 nm in aqueous suspensions, en, 2025.

(55) Hu, F.; Shi, L.; Min, W. Nature Methods 2019, 16, 830–842.

(56) Gentleman, E.; Swain, R. J.; Evans, N. D.; Boonrungsiman, S.; Jell, G.; Ball, M. D.; Shean, T. A. V.; Oyen, M. L.; Porter, A.; Stevens, M. M. Nature Materials 2009, 8, 763–770.

(57) Boonrungsiman, S.; Gentleman, E.; Carzaniga, R.; Evans, N. D.; McComb, D. W.; Porter, A. E.; Stevens, M. M. Proceedings of the National Academy of Sciences 2012, 109, 14170–14175.

(58) Napolitano, A. P.; Dean, D. M.; Man, A. J.; Youssef, J.; Ho, D. N.; Rago, A. P.; Lech, M. P.; Morgan, J. R. BioTechniques 2007, 43, 494–500.

