## Supplementary info for "Label-free imaging of matrix mineralization in alginate-encapsulated bone spheroids using Coherent Raman Scattering microscopy"

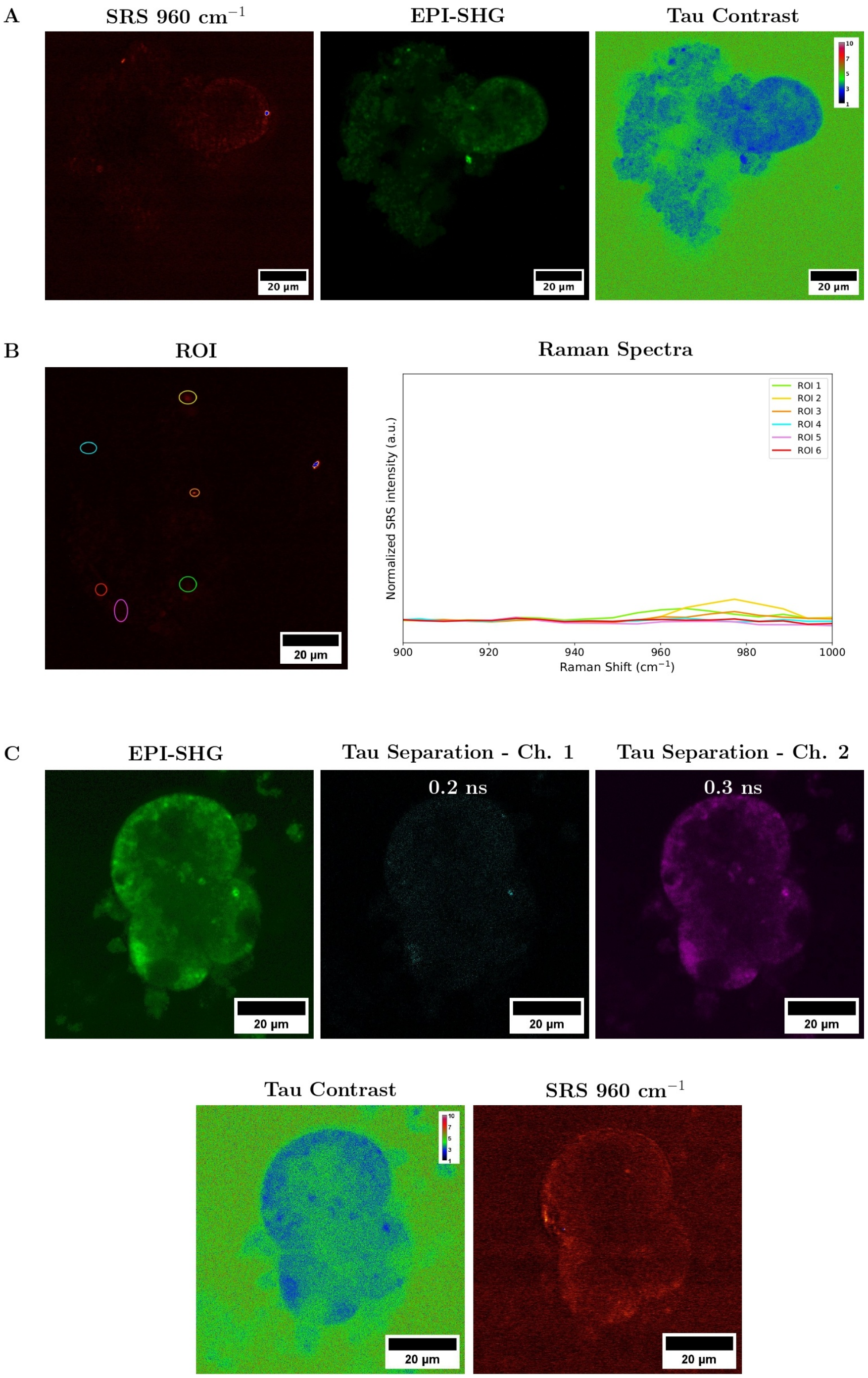


Figure S1: SRS and SHG analysis of mineralized ECM in alginate-encapsulated bone spheroids cultured for 4 weeks in RM. A) Combined SRS and epi-SHG images of a representative spheroid. The SRS image was obtained at Raman shift of 960 cm$^{-1}$ to detect phosphate groups associated with mineralized deposits. It can be observed that no signal was detected, indicating the absence of phosphate groups. The epi-SHG signal consist of contributions from the collagenous matrix and cellular auto-fluorescence; the signals were distinguished using the TauContrast function. The calibration bar represents the average arrival time in nanoseconds. Absence of collagen signal can be observed. B) Raman spectra acquired in the 900 to 1000 cm$^{-1}$ range and corresponding SRS image, indicating the selected regions of interest (ROIs). C) Epi-SHG images showing the signal contributions and the corresponding Tau-Separated channels, allowing the discrimination of collagen-derived SHG and auto-fluorescence based signal based on effective photon lifetime. No signal was detected for the fast channel associated with SHG signal, indicating the absence of deposited collagen. The TauContrast image and SRS image at 960 cm$^{-1}$ were collected simultaneously. The calibration bar in the Tau Contrast image represents the average arrival time in nanoseconds.

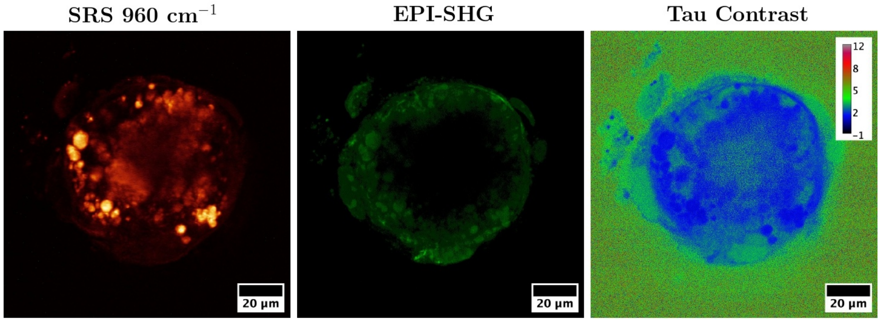


Figure S2: Validation of label-free SRS imaging through comparison with calcein fluorescence. Combined SRS, two-photon fluorescence and TauContrast images. Partial overlap was observed between SRS-positive regions in the outer layers of the spheroids and calcein-positive areas, indicating co-localization of phosphate-rich deposits with calcium-labeled structures. TauContrast was used to identify any change in the lifetime of the calcein-stained regions, compared to unstained samples.
